# Engineering of Co-Surfactant-Free Bioactive Protein Nanosheets for the Stabilisation of Bioemulsions Enabling Adherent Cell Expansion

**DOI:** 10.1101/2022.10.30.514404

**Authors:** Alexandra Chrysanthou, Minerva Bosch-Fortea, Julien E. Gautrot

## Abstract

Bioemulsions are attractive platforms for the scalable expansion of adherent cells and stem cells. In these systems, cell adhesion is enabled by the assembly of protein nanosheets that display high interfacial shear moduli and elasticity. However, to date, most successful systems reported to support cell adhesion to liquid substrates have been based on co-assemblies of protein and reactive co-surfactants, which limit the translation of bioemulsions. In this report, we describe the design of protein nanosheets based on two globular proteins, bovine serum albumin (BSA) and β-lactoglobulin (BLG), biofunctionalised with RGDSP peptides to enable cell adhesion. The interfacial mechanics of BSA and BLG assemblies at fluorinated liquid-water interfaces is studied by interfacial shear rheology, with and without co-surfactant acyl chloride. Conformational changes associated with globular protein assembly are studied by circular dichroism and protein densities at fluorinated interfaces are evaluated via surface plasmon resonance. Biofunctionalisation mediated by sulfo-succinimidyl 4-(N-maleimidomethyl) cyclohexane-1-carboxylate (sulfo-SMCC) is studied by fluorescence microscopy. On the basis of the relatively high elasticities observed in the case of BLG nanosheets, even in the absence of co-surfactant, the adhesion and proliferation of mesenchymal stem cells and human embryonic kidney (HEK) cells on bioemulsions stabilized by RGD-functionalized protein nanosheets is studied. To account for the high cell spreading and proliferation observed at these interfaces, despite initial moderate interfacial elasticities, the deposition of fibronectin fibers at the surface of corresponding microdroplets is characterized by immunostaining and confocal microscopy. These results demonstrate the feasibility of achieving high cell proliferation on bioemulsions with protein nanosheets assembled without co-surfactants and establish strategies for rational design of scaffolding proteins enabling the stabilization of interfaces with strong shear mechanics and elasticity, as well as bioactive and cell adhesive properties. Such protein nanosheets and bioemulsions are proposed to enable the development of new generations of bioreactors for the scale up of cell manufacturing.

## 1. Introduction

Bioemulsions, emulsions that are bioactive and support the culture of adherent cells, are emerging as attractive solutions for the scale up of cell manufacturing ^1,2^. Indeed, the absence of solid substrates or microcarriers is attractive to facilitate cell processing, reduce the contamination of cell products by microplastics and costs of culture consumables. Keese and Giaever first reported the culture of adherent cells at liquid interfaces, and identified the importance of surfactant molecules in mediating such process ^3,4^. More recently, the importance of these surfactants, or co/pro-surfactants, together with the self-assembly of proteins, on the determination of mechanical properties of associated interfaces was identified. Co-assembly of proteins or polymers, such as poly(L-lysine) (PLL), with reactive surfactant acyl chlorides led to the stiffening of interfacial mechanics at corresponding liquid-liquid interfaces ^5,6^. In turn, the resulting mechanically strong liquid-liquid interfaces are able to sustain shear forces developed by cells during their spreading and motility. In particular, the elastic properties of the protein assemblies at these interfaces, protein nanosheets, correlate with the proliferation of adherent stem cells such as mesenchymal stem cells and keratinocytes ^5,7,8^.

In this context, identifying assemblies that do not require co/pro-surfactants in order to strengthen interfacial mechanics is attractive, as it may enable faster translation of bioemulsions for stem cell manufacturing. Although a few systems have been proposed to date to promote stem cell adhesion and proliferation at liquid-liquid interfaces, they either rely on interfaces stabilized by undefined assemblies from serum proteins ^9^ or proteins that promote cell adhesion but do not readily stabilize emulsions, such as fibronectin ^10,11^. Therefore, the identification of scaffolding proteins that display tensioactive properties enabling the stabilization of emulsion, confer strong interfacial mechanics and present bioactive and/or cell adhesive ligands enabling cell adhesion and proliferation remains elusive.

Recently, the modification of globulins such as albumins with charged residues was found to impact their assembly at liquid-liquid interfaces and resulting nanosheet mechanics ^12^. These supercharged protein nanosheets enabled the simple adsorption of cell-adhesive proteins such as fibronectin and collagen, but the strengthening of their mechanical properties still required introduction of PFBC in order to sustain stem cell proliferation. However, this study demonstrated that simple protein modification may allow to modulate both nanosheet mechanics and bioactivity.

Globular proteins such as various globulins are attractive scaffold proteins for the design of nanosheets stabilizing bioemulsions as they can be sourced readily and are approved in a number of cases, at least for food applications, if not for use in therapeutics formulation. The functional properties of whey proteins are of substantial and growing importance to the food industry ^13^. Protein such as β-lactoglobulin are abundant, natural emulsifiers with relatively low cost. Associated food colloids are heterogeneous systems consisting of various kinds of particles and polymers. The nature and strength of interactions among them are the main determinants for the colloidal system properties that are highly influenced by the structure and composition of associated interfaces ^14^.

Important physicochemical properties that define the ability of a protein to form and stabilize emulsions are size, solubility, hydrophobicity, charge and flexibility ^15^. Strong viscoelastic films can provide electrostatic and steric stabilization, depending on the solvent conditions and the characteristics of the corresponding proteins. Flexible proteins such as caseins tend to form weaker viscoelastic films compared to globular compact proteins such as β-lactoglobulin, and may be more rapidly displaced by other surface active components ^15^.

During the formation of protein nanosheets at liquid interfaces, proteins must first travel from the bulk phase towards the interface via diffusion. Once at the interface, protein molecules then unfold in order to expose hydrophobic amino acids to the surface. This partial denaturation of the protein will mediate the rearrangement of the hydrophobic amino acids to face the oil phase, whilst hydrophilic amino acids rearrange towards the aqueous phase. The reorientation and unfolding of the hydrophilic and hydrophobic residues leads to the minimization of thermodynamic energy ^16^. The extent of conformational change depends on protein structure and the solvent conditions. Flexible proteins can rapidly adsorb at the interface, accelerating the reduction of interfacial tension without forming a dense and ordered packing layer. In contrast, globular proteins will pack at the interface, adsorbing at a slower rate but with higher order ^15^. In addition, during the adsorption, disulfide bonds can be formed, further contributing to emulsion stability ^17^.

β-lactoglobulin has a molecular weight of 18,500 Da, contains 5 cysteins forming 2 disulphide linkages and is rich in β-sheet structures ^18,19^. β-lactoglobulin consists of three-turn α-helix and two β-sheets made by nine strands that are folded forming a hydrophobic calyx, classifying β-lactoglobulin into the lipocalin proteins family ^20^. This formed calyx makes β-lactoglobulin able to bind to hydrophobic vitamins or lipids ^21^. β-lactoglobulin has an internal free sulfhydryl group which is only available as the protein adsorbs and partially unfolds ^13^. During adsorption at the interface, β-lactoglobulin was proposed to form intermolecular β-sheet leading to the development of a strong protein viscoelastic nanosheet at the interface ^15^. The high viscoelasticity of this film is further supported by intermolecular disulfide bonds due to the free thiol group present in the structure ^15^. The high ordered packing thus created, together with intermolecular crosslinks, makes the disruption of the film by other molecules such as surfactants extremely difficult ^15^.

Therefore, β-lactoglobulin appears as an attractive scaffolding protein candidate for the design of viscoelastic protein nanosheets able to resist cell-mediated traction forces generated during spreading and proliferation. However, the lack of available integrin ligands able to initiate processes resulting in cell spreading requires further design of this protein. In addition, further combination with co-surfactant molecules such as acyl chlorides has not been studied.

Various approaches have been proposed to functionalize globular proteins with peptides. For example, bovine serum albumin (BSA) was functionalized with cyclic-RGD presenting free cysteines at the carboxylic end in order to mediate chemical coupling to maleimide residues introduced on the albumin through the heterobifunctional reagent sulfo-succinimidyl 4-(N-maleimidomethyl) cyclohexane-1-carboxylate (sulfo-SMCC) ^22^. This enabled to mediate cell adhesion to resulting albumin films that otherwise would block cell spreading ^22^.

In this report, the viscoelastic behavior of BSA and β-lactoglobulin adsorbed at fluorinated oil interfaces are first investigated via interfacial shear rheology. The impact of the co-surfactant pentafluorobenzoyl chloride (PFBC) on this process is then studied. Conformational changes associated with the adsorption of these proteins at fluorinated oil interfaces are characterized via circular dichroism. The functionalization of resulting protein nanosheets with RGD cell adhesive peptides, using sulfo-SMCC as coupling agent, is then investigated. Finally, the proliferation of two adherent cell types, mesenchymal stem cells used in stem cell therapies, and HEK293 cells used for the expression of recombinant proteins, at the surface of bioemulsions stabilized by RGD-functionalized BSA and β-lactoglobulin is investigated. Cell adhesion and cytoskeleton assembly at the surface of nanosheet stabilized oil droplets are characterized and extra-cellular matrix deposition is studied.

## 2. Results and Discussion

The formation of protein nanosheets based on bovine serum albumin (BSA) and β-lactoglobulin (BLG) at oil-water interfaces was first characterized, using interfacial rheology (Fig. 1). Focus was placed on interfaces formed with the fluorinated oil Novec 7500, due to its broad application in a range of microdroplet microfluidic technologies and its low cytotoxicity^23,24^. Upon injection of BSA and BLG solutions, the interfacial shear storage modulus of corresponding liquid-liquid interfaces increased by two to three orders of magnitude, as shown in the Fig. 1A. This is in agreement with previous reports indicating that albumin readily adsorbs to a range of oil-water interfaces, forming a highly viscoelastic interface with interfacial dilatational storage^25–28^ moduli in the range of 20–60 mN/m, depending on the technique used. The variation in values measured likely reflects variations in batch and source of the proteins used, but also the type of interfacial rheology applied. In particular, techniques relying on droplet shape analysis result in moduli that depend on surface tension, as well as the shear moduli of corresponding interfaces, rather than purely on shear properties ^29^. Indeed, surface tensions are comparable to dilatational interfacial shear moduli of many protein-stabilized liquid-liquid or liquid-air interfaces ^29,30^, and contribute to significant levels to interfacial mechanics, in addition to coulombic interactions, assessed by dilatational rheology and via AFM indentations ^31^. β-lactoglobulin led to the formation of interfaces with comparable interfacial storage moduli, compared to BSA (Fig. 1A and B). These data are in good agreement with the relatively fast adsorption of β-lactoglobulin and BSA at other hydrophobic liquid interfaces, whilst adsorption to more polar hydrophobic liquid was retarded and led to weaker interfacial mechanics ^30,32,33^.

**Figure 1.**
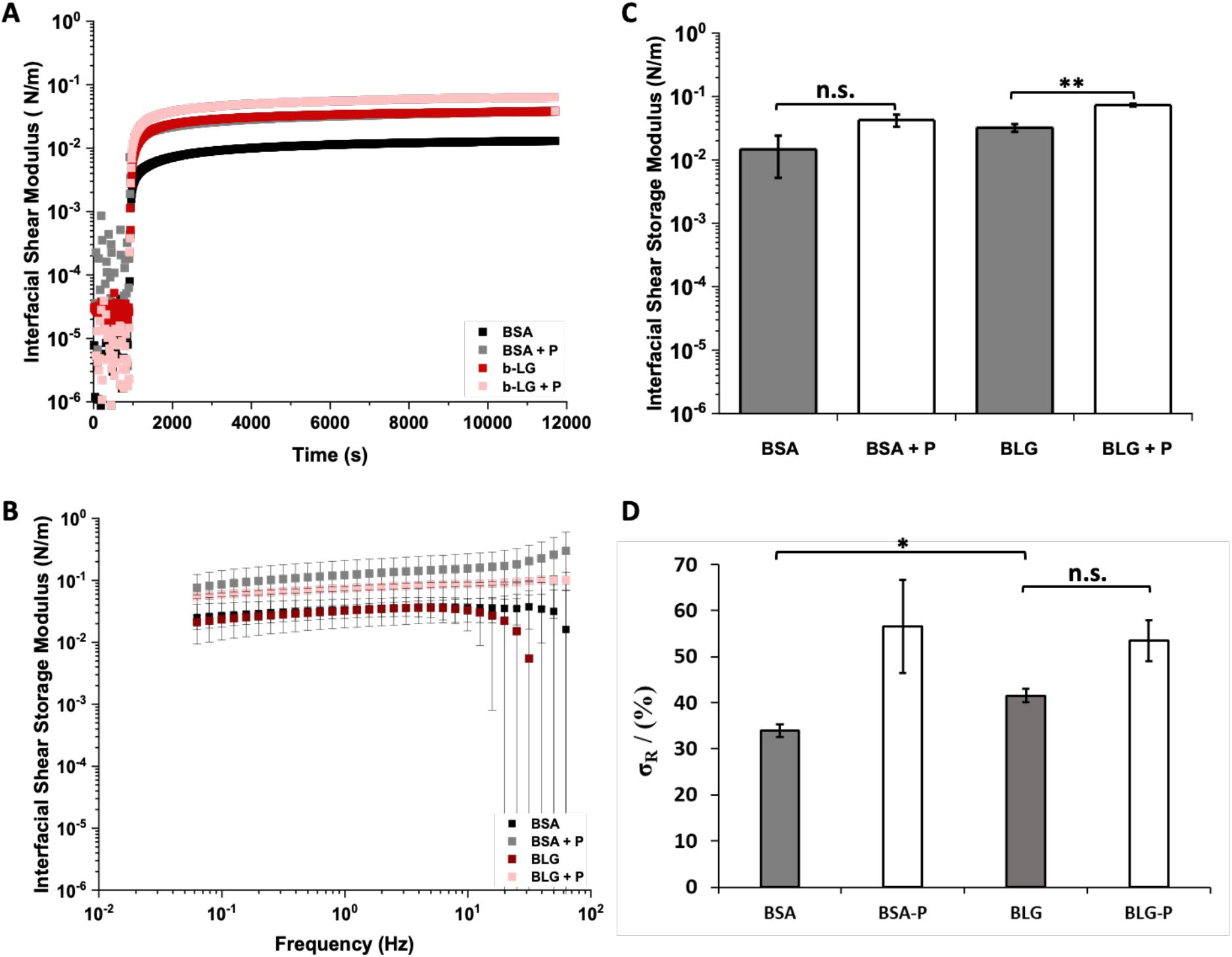
A) Evolution of interfacial storage moduli during the adsorption of BSA and β-lactoglobulin, with and without PFBC (10 μg/mL with respect to the oil phase). B) Frequency sweep of the protein nanosheets and C) the corresponding interfacial storage modulus at oscillating amplitude of 10^−4^ rad and frequency of 1 Hz. D) Residual interfacial elasticities (stress retentions, σ_R_, %) extracted from the fits of stress relaxation experiments at 0.5 % strain (see Supplementary Figure S2 for representative examples of traces).

The interfacial shear moduli of the generated interfaces was found to be moderately dependent on oscillating frequencies, indicating the formation of viscoelastic interfaces (Fig. 1C). In addition, frequency sweeps further confirmed that β-lactoglobulin formed interfaces with equivalent interfacial shear moduli to BSA. These results were further confirmed by comparison of interfacial storage and loss moduli data (Fig. 1B and Supplementary Fig. S1). The interfacial loss moduli of BSA and BLG were both nearly one order of magnitude lower than corresponding interfacial storage moduli. However, whilst the interfacial shear storage modulus of BLG was found to be higher than that of BSA, its interfacial loss modulus was not. To further investigate viscoelastic properties of BSA and BLG interfaces, interfacial stress relaxation experiments were carried out using recently established protocols^8^ (Supplementary Fig. S2). Significant levels of elasticity (high elastic stress retention, σ_R_) were observed in BLG-stabilized interfaces, compared to BSA. Therefore, these data indicate that whilst BSA and BLG interfaces display comparable interfacial storage moduli, their elasticity differs, suggesting more extended crosslinked networks are obtained for BLG.

The addition of co-surfactant molecules was previously found to significantly affect interfacial mechanics, in particular viscoelastic profiles^5,7,8^. Therefore, the impact of introduction of the pro-surfactant pentalfluorobenzoyl chloride (PFBC) was studied next (Fig. 1A). Acyl chlorides such as PFBC are referred to as co-surfactants, rather than surfactants, as they display moderate tensioactive properties, and instead couple covalently to proteins or polymers to promote their adsorption or modulate the mechanics of resulting interfaces ^5,7,8^. Both BSA and BLG interfaces were found to display higher storage and loss moduli in the presence of PFBC (Fig. 1A-C and Supplementary Fig. 1), indicating that coupling of these hydrophobic residues enhanced physical crosslinks within the protein layer. Indeed, a wider range of hydrophobic acyl chloride was previously reported to induce nanosheet strengthening through the formation of hydrophobic crosslinks^8^. Changes in interfacial viscoelastic behavior were particularly striking, with enhancement in elastic stress from 34 and 42 % for BSA and BLG interfaces (respectively) to 57 and 54 % for BSA and BLG nanosheets formed in the presence of PFBC (Fig. 1D and supplementary Fig. S2). Considering the higher elasticity observed for BLG interfaces in the absence of PFBC, the magnitude of the change in stress retention observed in relaxation experiments was found to be stronger for BSA nanosheets. This difference may also stem from the significantly higher number of lysine residues (near 60 per molecule) in BSA compared to β-lactoglobulin (only 15 lysines). As the amines of lysines are proposed to couple to acyl chlorides (although other residues such as serines may also contribute to reactivity), the presence of higher lysine densities may enhance physical crosslinking and associated elasticity of BSA nanosheets. The formation of highly elastic protein nanosheets, in the absence of co-surfactant is attractive to enable faster translation of these assemblies, due to the lack of validation of these molecules, including PFBC, by regulatory bodies. Therefore, BLG is an attractive candidate for the stabilization of liquid-liquid interfaces and strengthening of their mechanical properties in the absence of PFBC or other co-surfactants, to enable cell adhesion and proliferation on liquid substrates such as microdroplets. Although the impact of BLG on surface tension, the stabilization of emulsions and interfacial shear mechanics is well established^34–36^, the origin of its strengthening of interfacial shear mechanics is incompletely understood.

To explore the mechanism associated with β-lactoglobulin strengthening of interfacial shear mechanics, the conformation of BSA and BLG was investigated in solution and at liquid-liquid interfaces. β-lactoglobulin consists in 33% β-sheet structures, which is significantly higher than the β-sheet content of BSA (10%) ^37–39^. This high percentage of β-sheet confers to BLG a β-structure that is more rigid than other α-helix-dominated or disordered proteins ^40–42^. The higher rigidity of β-sheet-rich proteins was studied Boson Peak Frequency by Perticalori et al., concluding on the increased stiffness of β-structured proteins compare to that of α-structured proteins^43^. Upon adsorption at hydrophobic liquid interfaces, rearrangement of protein structure may significantly impact on interfacial mechanics by enabling exposure of hydrophobic residues that may provide intermolecular crosslinks ^32,44^. In addition to structural changes, free thiol groups may contribute to further crosslinking of associated protein networks and modulate interfacial mechanics or stabilize emulsions ^13,45,46^.

To study conformation changes in BSA and BLG, circular dichroism (CD) measurements were carried out. In order to enable such measurements, mixtures of Novec 7500 and α,α,α-trifluorotoluene (1.5:1 ratio) were used (refractive index of Novec 7500 and of α,α,α-trifluorotoluene –– 1.29 and 1.41, respectively) ^47^, with refractive index matching that of PBS buffer. This afforded clear emulsions that enabled CD measurements (Fig. 2A). The CD spectra of BSA and BLG solutions was in good agreement with their expected structure and reports from the literature ^48,49^ (Fig. 2B/C). BLG, in solution, was found to exhibit strong beta-sheet components and associated profile, with the characteristic positive maximum at 195 nm, the negative maximum at 218 nm and the zero crossings at 207 and 250 nm ^49^. To quantify the conformational contribution associated with such spectra, the SELCON algorithm from Dichroweb was used ^50^. The calculated composition of BLG structures in solution was found to be 17% α-helix, 41% β-sheet and the remaining corresponding to disordered and β-turns structures, consistent with previous results and the estimated secondary structure (Table 1) ^49^. In contrast, in solution, BSA displayed strong α-helical components, with a positive peak at 195 nm, a double negative peak at 210 and 222 nm, and a zero crossing at 205 nm (Fig. 2C). This corresponded to a α-helix content of 72%, with no significant β-sheet contribution and the rest accounted by β-turns and disordered domains, in good agreement with the expected structure of BSA and previously reported CD data ^19,38^.

**Figure 2.**
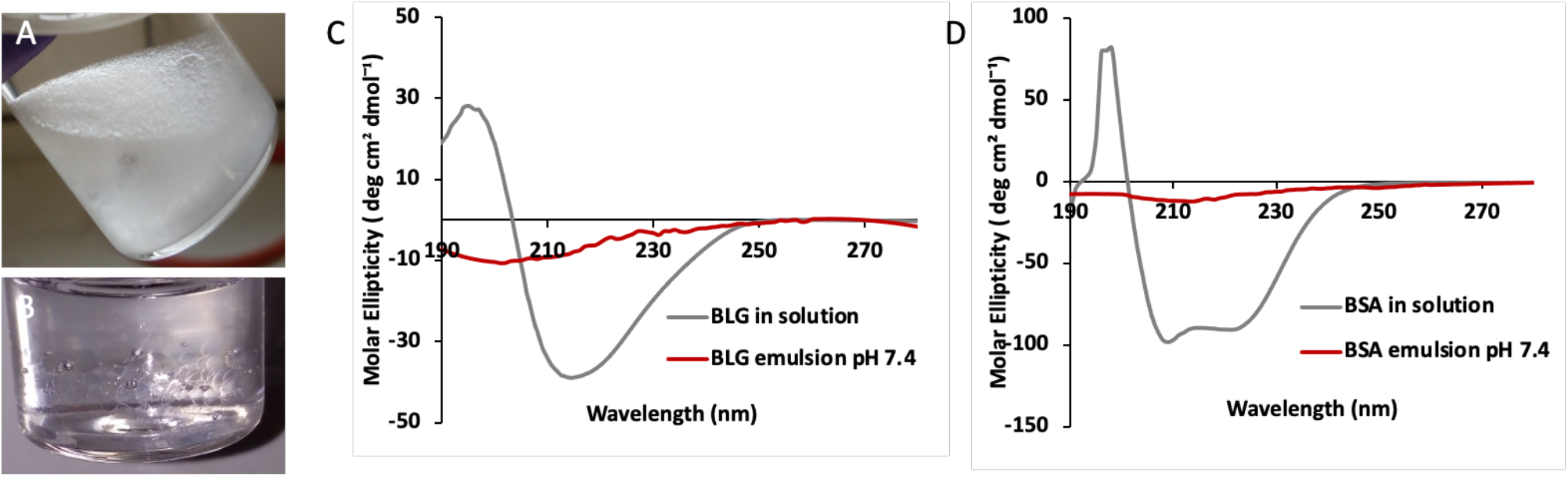
A) Representative image of a Novec 7500-water emulsion. B) Refractive index matched-emulsion made with a mixture of Novec 7500 and trifluorotoluene. CD spectra of C) BLG in solution (at concentration 1 mg/ mL) and adsorbed in an oil-water interface (after removing excess free protein) and D) BSA in solution (at concentration 1 mg/ mL) and adsorbed at oil-water interface (after removing excess free protein).

**Table 1.** Secondary structure composition of BSA and BLG in solution and at interfaces calculated using SELCON algorithm in DichroWeb, based on circular dichroism data.

|  | Secondary Structure (%) |  |  |  |
| --- | --- | --- | --- | --- |
|  | Helix | Sheet | Turns | Unordered |
| **BSA in solution** | 72 | 0 | 10 | 18 |
| **BSA in emulsion** | 25 | 13 | 14 | 48 |
| **BLG in solution** | 17 | 42 | 26 | 15 |
| **BLG in emulsion** | 17 | 31 | 12 | 39 |

Significant changes in CD spectra were observed in emulsions (Fig. 2B/C). The α-helix contribution of BSA reduced to only 26%, whereas some β-sheet contribution occurred (13%) and the disordered contribution increased to 48%. In contrast, the α-helix composition of BLG remained unchanged and its β-sheet component increased to 31%, with an increase of disordered domains (from 15 to 38%). Hence, CD spectra indicate a significant unfolding of BSA and BLG at fluorinated oil interfaces, with a predominant disordered structure. Similar trends were observed for BSA and myoglobin adsorbed at hexadecane/water interface showing an increase of disordered structure (17 to 27% and 13 to 29%, respectively) ^19^. It should be noted that molar ellipticities were based on the starting concentration of protein in solution (in the aqueous phase) and that concentrations at the interfaces of emulsions were not corrected, despite the aqueous phase having been washed and exchanged for protein free buffer.

These results are in general agreement with previous reports, in which FT-IR spectra indicated high β-sheet contents for BLG in solutions (45%), with minor α-helix content (7%) ^51^. Upon adsorption to triacyglycerol (TAG), β-sheet components were found to reduce modestly to 25% while, α-helix and random coil contributions increased from 7% to 25% and from 20% to 27%, respectively ^52^. Similarly, β-sheet content of BLG adsorbed at diacyglycerol (DAG)-water interfaces was reduced from 45% to 35%, although a moderate level compared to BLG at TAG interfaces, while the α-helix component increased from 7% to 20%^52^. It was suggested that the enhanced unfolding at DAG-water interfaces, compare to TAG-water, results from the associated reduced surface coverage. Such decrease in surface coverage was proposed to originate from the higher polarity of the DAG compare to TAG^51^. Indeed, surface coverage correlated with the polarity of TAG and n-tetradecane ^51^. These changes were also comparable to those reported by Zhai et al., indicting partial loss in β-sheet and an increase in α-helix composition and disordered domains^40,53^. Other studies have also proposed the conversion of α-helical rich proteins such as BSA, lysozyme and myoglobin to β-sheet components upon adsorption to oil-water interfaces ^19^. Therefore, overall, our results suggest a significant change in protein structure upon assembly at fluorinated oil interfaces. Therefore, it is possible that intermolecular interactions between residues exposed upon such conformational rearrangement may underlie physical crosslinking of associated protein nanosheets and the level of elasticity observed, even in the absence of PFBC coupling. However, further crosslinking seems to require additional residues, introduced through PFBC molecules. The high hydrophobicity of the fluorinated oil studied may also contribute to enhance surface coverage and associated protein entanglement, facilitating crosslinking. Finally, the oligomerization state of BLG, known to be modulated by the pH of associated solutions, may also impact on interfacial assembly and mechanics, although at the pH of the present study, BLG was reported to be predominantly dimeric ^54–57^,.

Despite these encouraging interfacial mechanical properties, BSA and BLG remain inherently globular proteins playing roles in molecular transport and lipid stabilization in physiological fluids ^58–60^, with little relevance to ECM signaling and the promotion of cell adhesion. To confer cell adhesion to these scaffold proteins, we coupled an RGD peptide to the surface of protein nanosheet-stabilized droplets. The heterobifunctional coupling agent sulfo-SMCC was allowed to tether to nanosheets, prior to washing of excess and incubation in cysteine-terminated peptides displaying the cell adhesive ligand RGDSP ^22^. To determine the success of this tethering strategy, the thiolated dye 5-((S-(acetylmercapto)succinoyl amino fluorescein (SAMSA) was coupled instead of peptides, prior to quantification via fluorescence microscopy (Fig. 3). Michael-addition of SAMSA was carried out to sulfo-SMCC activated droplets stabilized by BSA and BLG nanosheets. This resulted in homogenous levels of functionalization, over the droplet surface and throughout the droplet population (Fig. 3). Quantification of the fluorescence indicated overall comparable levels of functionalization on BSA and BLG nanosheets, with excellent retention of emulsion stability (note that the emulsions studied were generated by vortexing and inherently polydisperse in size). The control groups, maleimide-free, did not display any significant functionalization or conjugation with SAMSA, demonstrating that the coupling is maleimide-specific and enabling the potential control of ligand density. The lack of fluorescence from pristine nanosheet-stabilized emulsions (without sulfo-SMCC or SAMSA) also confirmed the lack of auto-fluorescence from adsorbed proteins. Taken together, these results demonstrate the capability of this approach to directly functionalize a variety of protein nanosheet-stabilized microdroplets, a flexible approach to systematically combine peptide formulations.

**Figure 3.**
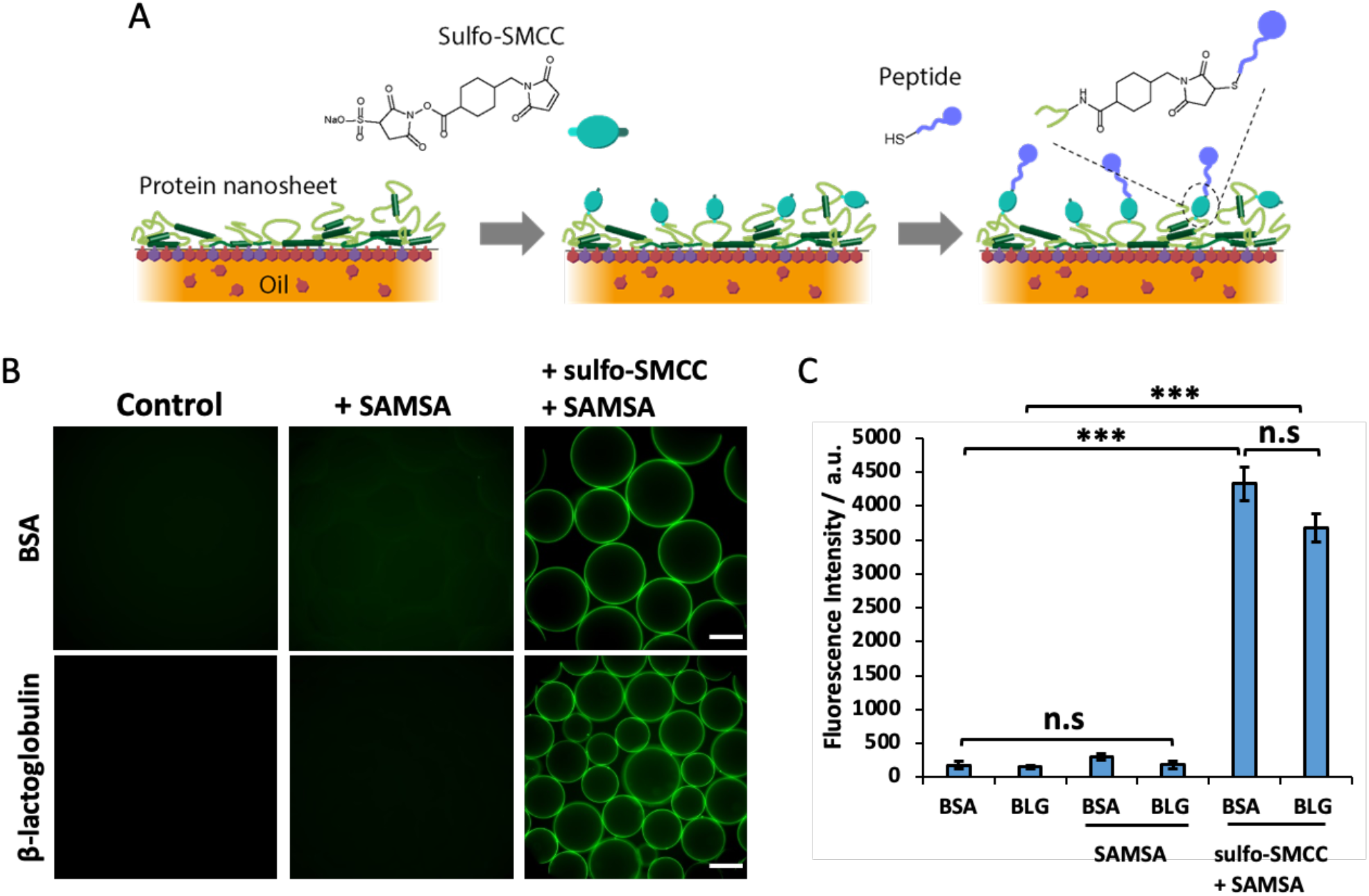
A) Schematic representation of the functionalisation protocol. B) Epifluorescence microscopy images of BSA or BLG emulsions functionalised with SAMSA fluorescein (green, A685) with or without sulfo-SMCC. C) Mean fluorescence intensity of the coupled SAMSA. Scale bars, 100 μm. Error bars are s.e.m.; n = 3.

To further investigate the functionalization process, surface plasmon resonance (SPR) was used to characterize associated successive steps (Fig. 4). To better capture the structure of nanosheets formed at fluorinated liquid interfaces, SPR chips were functionalized with a monolayer of perfluorinated thiol. This is potentially leading to adsorption mechanisms that will differ from adsorption to Novec 7500, but would better capture representative adsorption/functionalization profiles to fluorinated oils than direct adsorption to gold surfaces (of the SPR chips). The adsorption of BSA and β-lactoglobulin was found to be comparable (Fig. A-B), although perhaps associated with slightly thicker or denser BLG assemblies. The adsorption levels measured for BSA (120 ng/cm^2^) and BLG (180 ng/cm^2^) were not statistically different and corresponded to protein sub-monolayers in both cases. Indeed, perfectly packed monolayers of BSA would give rise to densities ranging from 190 to 630 ng/cm^2^ (depending on the orientation and assuming no unfolding; taking dimensions of 141 × 42 × 42 nm), whereas BLG would give rise to densities near 230 ng/cm^2^ (assuming a closely packed sphere of 3.6 nm diameter). Therefore the adsorption levels observed suggest either sub-monolayer formation, or significant unfolding. In turn, coupling of sulfo-SMCC was associated with changes in surface densities of 70 and 80 ng/cm^2^ for BSA and BLG respectively, presumably corresponding to the mass increase associated with coupling and potential further changes in protein conformation (see Fig. 4 C-D). Finally, peptide coupling was associated with further increase in mass densities of 14 and 16 ng/cm^2^ (for BSA and BLG, respectively; comparable for both proteins; see Fig. 4 E-F). The kinetics at which the reaction took place was slower, reflecting the lower concentration of peptides compared to sulfo-SMCC, and despite the higher molar mass of the cell adhesive peptide selected. However, with a molar mass of 805 g/mol, this level of adsorption remains associated with a degree of coupling of 12 and 2 peptides per protein (BSA and BLG, respectively) and peptide surface densities of 0.10 and 0.12 peptides/nm^2^. Therefore, the distance between each peptide for both bioactive protein nanosheet interfaces is below 10 nm, clearly under the threshold beyond which cells are able to sense reductions in adhesive ligand densities ^61^. Therefore, the maleimide-based coupling of RGD peptides to BSA and BLG nanosheets was found to be well within the densities that are considered suitable to enable rapid cell adhesion and spreading.

**Figure 4.**
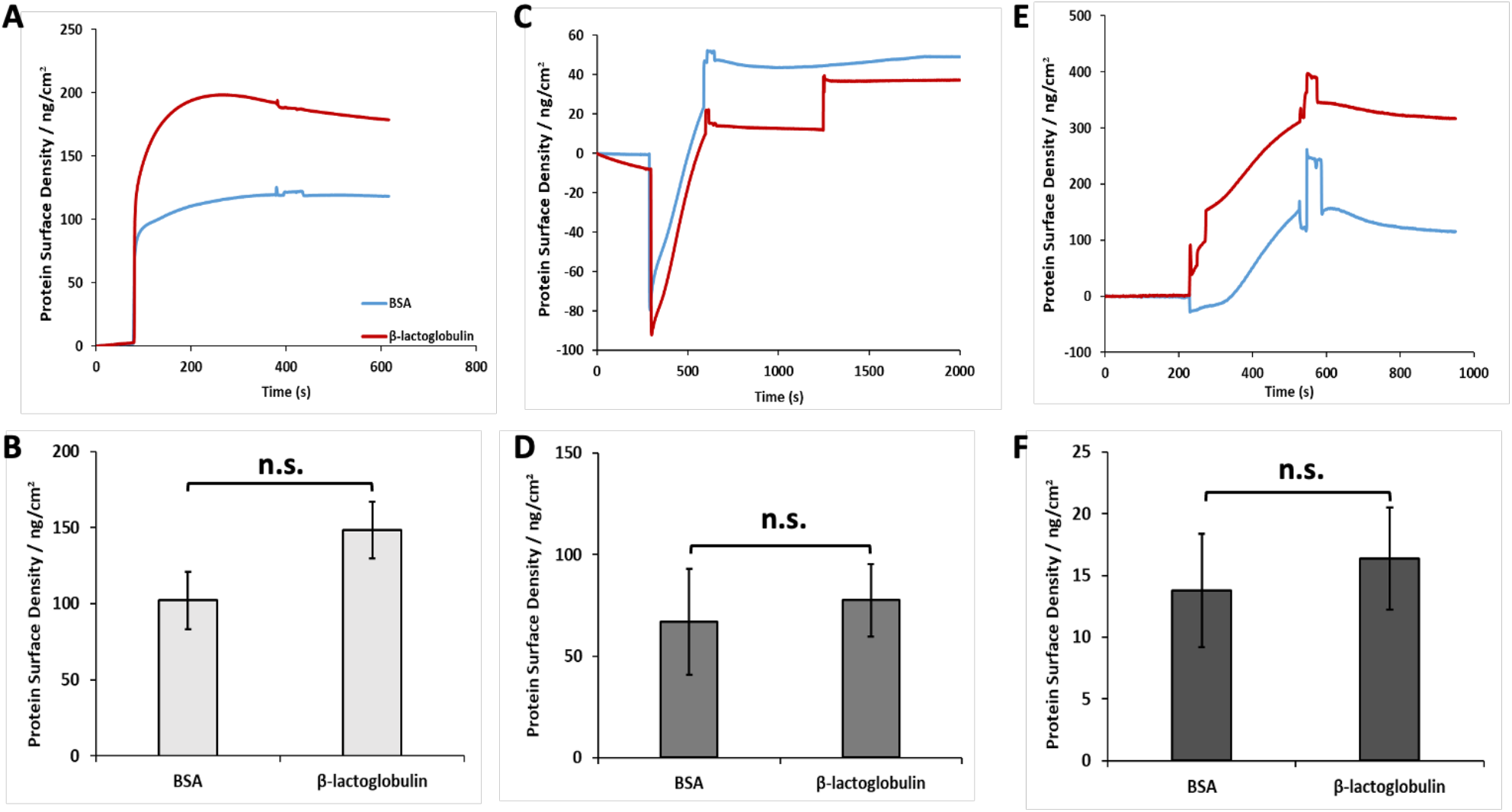
A) Representative surface plasmon resonance traces displaying the adsorption of BSA and BLG to perflurodecanethiol monolayers modelling fluorinated oil interfaces. B) Corresponding quantification of resulting protein surface densities. C) SPR quantification of sulfo-SMCC coupling at the surface of BSA and BLG layers. D) Corresponding calculated additional mass. E) SPR quantification of RGD coupling at the surface of sulfo-SMCC layers. F) Corresponding peptide surface densities. Error bars are s.e.m.; n = 3.

Having demonstrated the suitable biofunctionalisation of protein nanosheets with cell-adhesive ligands, the culture of mesenchymal stem cells (MSCs) at these interfaces was next investigated. MSCs were cultured at the surface of oil droplets stabilized by RGD-functionalized protein nanosheets. As a comparison, in addition to tissue culture plastic (TCP) controls, poly(L-lysine) (PLL) nanosheet-stabilized emulsions were also investigated, as the expansion of MSCs to such interfaces has previously been reported and characterized extensively^5^. After 7 days of culture, MSCs were found to have proliferated significantly at the surface of BSA and BLG nanosheet-stabilized emulsions functionalized with RGD peptides, to levels comparable to those observed on TCP and PLL-stabilized microdroplets (Fig. 5A). Cell colonies were found to cover droplets relatively homogenously, with high droplet occupancies (Fig. 5B and Supplementary Fig. S5). At day 7, cell densities were slightly higher on BLG/RGD-emulsions than BSA/RGD-emulsions. Surprisingly, although very few cells were found on BSA-emulsions, cell densities at the surface of BLG-emulsions were comparable to those observed on BSA/RGD-emulsions and only slightly below those of BLG/RGD-emulsions. As no cell-adhesive ligand has been reported within the structure of BLG, to the best of our knowledge, these results suggest instead that matrix adsorption to BLG dominates this behavior. Interestingly, although BSA nanosheets were found to display relatively low elasticities, cell adhesion and proliferation at these interfaces was found to be relatively high, without supplementation with PFBC. This may suggest further maturation of the mechanical properties of corresponding nanosheets, perhaps in response to matrix adsorption.

**Figure 5.**
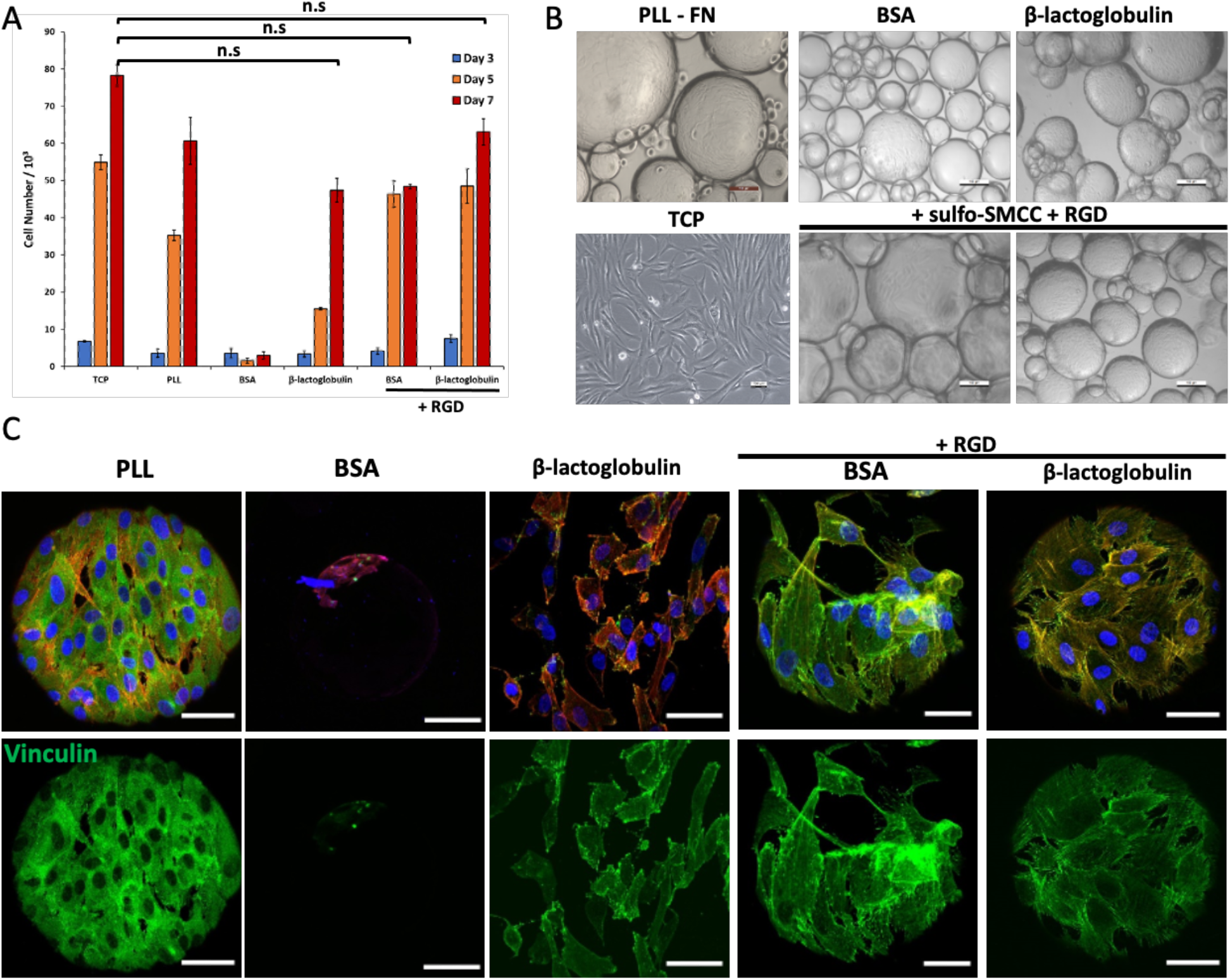
A) Proliferation of mesenchymal stem cells (MSC) at the surface of bioemulsions stabilized PLL, BSA or BLG nanosheets, with or without RGD functionalization. Based on metabolic assay. Comparison with tissue culture plastic (TCP) control. B) Corresponding bright field images of MSCs cultured for seven days. C) Confocal images of MSCs cultured for seven days on corresponding bioemulsions and controls (blue, DAPI; red, phalloidin; green, vinculin;). Scale bars are 100 μm (bright field) and 50 μm (confocal). Error bars are s.e.m.; n = 3.

To further investigate cell adhesion to nanosheet-stabilized emulsions, the formation of focal adhesions and cytoskeleton assembly were investigated (Fig. 5C). Although quantification was not carried out, due to the difficulty of imaging such curved interfaces at high resolution, with suspended droplets, images clearly indicated the formation of focal adhesions and establishment of a mature cytoskeleton on BSA/RGD and BLG/RGD bioemulsions, comparable to the phenotype of MSCs spreading to PLL-stabilized emulsions. In contrast, the few cells that were found to spread on BSA-emulsions were relatively rounded. However, in agreement with the high proliferation of MSCs at BLG-emulsions, despite the lack of RGD functionalization, cells were found to form focal adhesions and assemble a cytoskeleton on BLG nanosheets, although their spreading seemed slightly reduced compared to BLG/RGD nanosheets. Therefore, these results indicate that the expansion of MSCs at the surface of BSA/RGD and BLG/RGD-stabilized bioemulsions is mediated by cell adhesive ligands and regulated by the classic acto-myosin machinery, as was previously observed on fibronectin coated PLL-stabilized liquid-liquid interfaces ^5^. These results are consistent with the high interfacial moduli and relatively high elasticities observed for these nanosheets, in particular BLG, as well as high ligand densities achieved, as both of these surface properties are classically associated with the regulation of cell adhesion, spreading and proliferation at various interfaces ^62–64^.

A surprising aspect of these results is that cell adhesion and spreading was found to be excellent on both BSA/RGD and BLG/RGD nanosheets, despite the moderate elasticities measured for both interfaces, in the absence of PFBC (Fig. 1), as the ability of liquid-liquid interfaces to store strain energy and resist deformation was found to be highly correlated with adherent cell expansion at liquid-liquid interfaces ^65^. To further explore some of the possible mechanisms via which cell adhesion to droplets stabilized by relatively viscous nanosheets (low interfacial elasticity), matrix deposition was investigated (Fig. 6). MSCs culture for 7 days on bioemulsions deposited fibronectin fibres that covered the surface of droplets and formed relatively dense networks. Interestingly, this was also the case at the surface of BLG-emulsions, with matrix deposition also visible in gaps within cell colonies, suggesting that matrix remodeling underpins at least some of the adhesion and proliferation of MSCs at these interfaces. In addition, on the few BSA-stabilized emulsions that supported MSC adhesion, some fibronectin assembly was clearly visible, suggesting that such phenomenon might contribute to the proliferation of these rare colonies. This may be associated with the direct assembly of fibronectin (or other ECM proteins) at hydrophobic liquid interfaces, as this process was found to be sufficient to support the adhesion and proliferation of some stem cells (although not on emulsions, given the poor tensioactive properties of fibronectin)^11^. Alternatively, it may be that fibronectin (or other ECM molecules) are able to adsorb to assembled denatured BSA.

**Figure 6.**
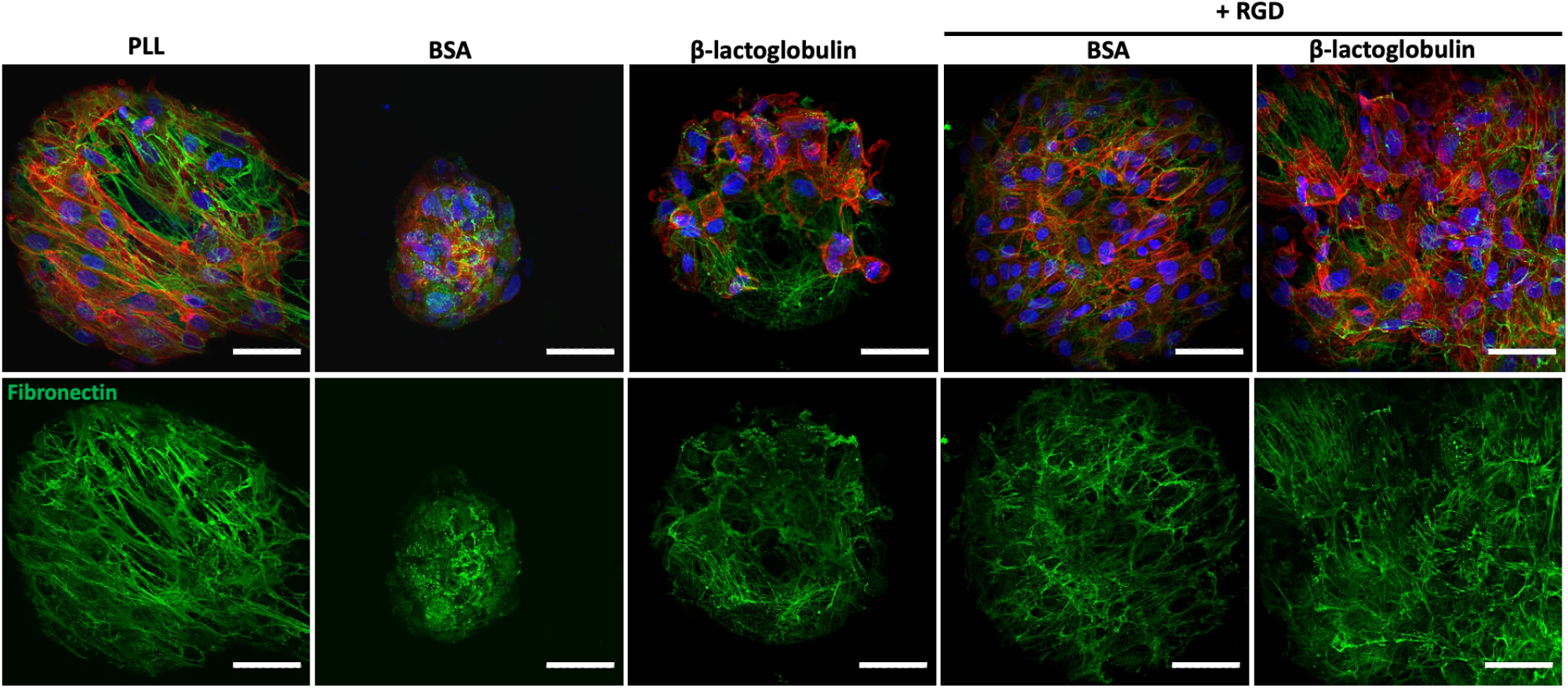
Confocal images of MSCs cultured for seven days on bioemulsions stabilized by protein nanosheets. Characterization of fibronectin deposition (blue, DAPI; red, phalloidin; green, fibronectin;). Scale bars are 50 μm (confocal). Error bars are s.e.m.; n = 3.

**Figure 7.**
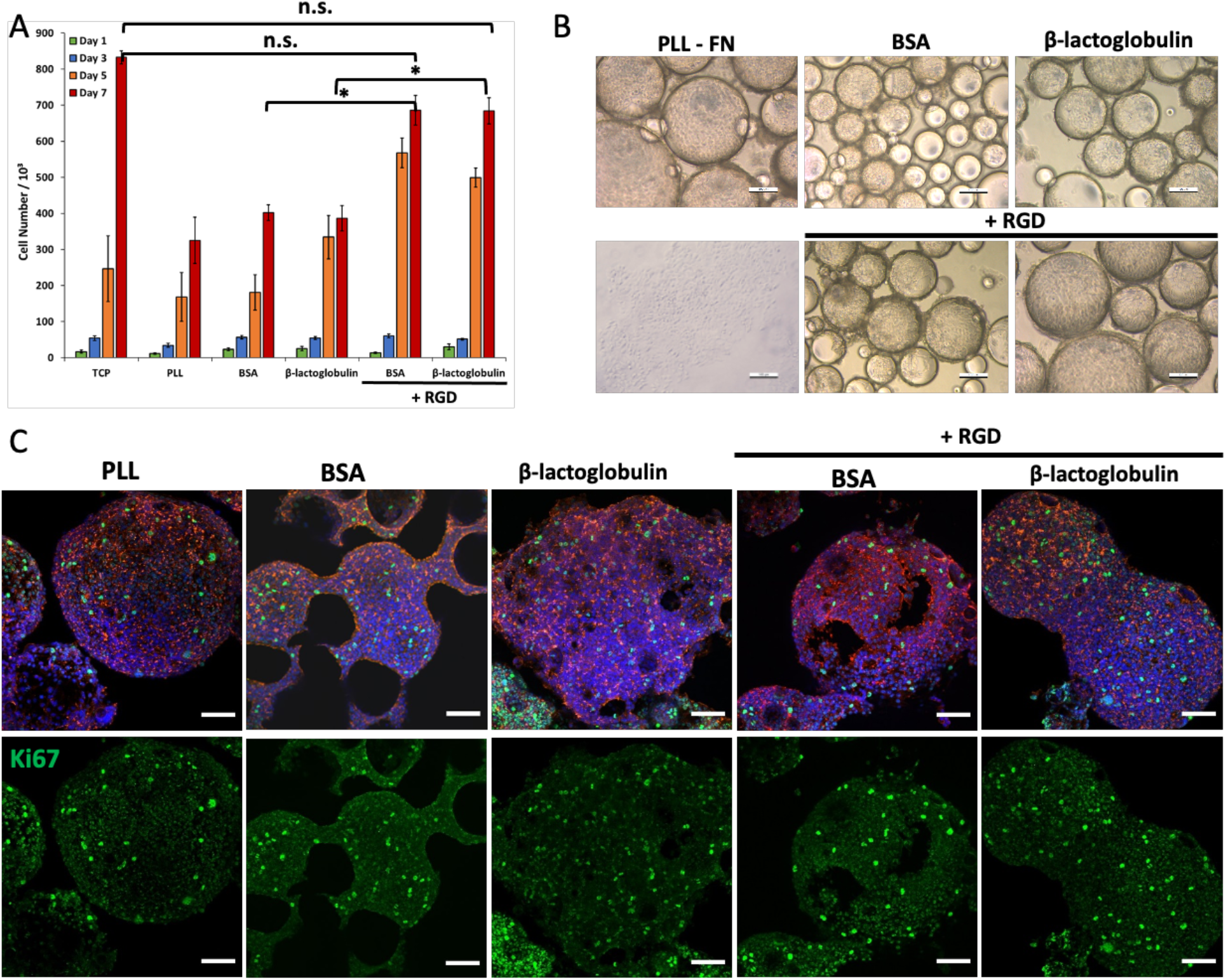
A) Proliferation of Human embryonic kidney cells (HEK293T) at the surface of bioemulsions stabilized PLL, BSA or BLG nanosheets, with or without RGD functionalization. Based on metabolic assay. Comparison with tissue culture plastic (TCP) control. B) Corresponding bright field images of MSCs cultured for seven days. C) Confocal images of MSCs cultured for seven days on corresponding bioemulsions and controls (blue, DAPI; red, phalloidin; green, vinculin;). Scale bars are 100 μm (bright field) and 50 μm (confocal). Error bars are s.e.m.; n = 3.

Finally, the culture of human embryonic kidney cells (HEK293T), often used for the expression of recombinant proteins by mammalian cells ^66,67^, was explored at the surface of bioemulsions, to demonstrate the potential of these platforms for the production of biotherapeutics and recombinant proteins. Cell densities were characterized at days 1, 3, 5 and 7 (Fig. 6). Interestingly, all conditions were found to support relatively high cell densities, compared to those observed for MSCs. However, HEK293 proliferation was particularly strong at the surface of BSA/RGD and BLG/RGD emulsions, comparable to that observed on TCP. In non-functionalized BSA and BLG emulsions, although relatively high proliferation was observed, many droplets could be seen to be devoid of colonies and HEK293 were found to form aggregates that seem weakly adhered to neighboring droplets. This may be associated with the capacity of HEK293 cells to sustain moderate proliferation in suspension, or in weak adhesive states ^68^. To confirm the high proliferation observed on these various interfaces, Ki67 staining, a marker reflecting cell cycling, was investigated (Fig. 6C). This indicated clear levels of proliferation on all emulsions, supporting the bright field imaging and cell density quantification data.

## 3. Conclusions

In this study, the possibility to grow cells on microdroplets stabilized by defined scaffold proteins, in the absence of further co-surfactant assembly, was demonstrated. Protein nanosheet assembly is found to be associated with significant denaturation and conformational rearrangement, depending on the protein type. Such conformational changes may be the basis for the formation of relatively elastic networks interconnecting assembled proteins and underpinned by re-association between rearranged residues, although such processes remains to be demonstrated formally. However, with β-lactoglobulin too, as previously shown with albumins^7^, the mechanics of these networks can further mature in the presence of PFBC. Overall, supramolecular interactions can drive the formation of quasi-2D elastic protein networks. Covalent strategies to further mature and further tune the mechanics of these networks are expected to bring further control to these interfaces, as was recently demonstrated in the case of simply exposure to DTT, likely promoting disulfide bond formation^69^.

In addition, this study demonstrates a straight forward strategy for the biofunctionalisation of pre-assembled scaffold proteins and corresponding emulsions, with cell adhesive peptides. Sulfo-SMCC mediated coupling can be readily applied to a variety of other scaffold proteins and a broad range of peptides sequences can be selectively coupled to these residues through Michael additions. Finally, this study demonstrates that the resulting bioactive nanosheets support the adhesion and proliferation of MSCs and HEK293 cells at the surface of corresponding bioemulsions. β-lactoglobulin-stabilized bioemulsions are found to perform better in this respect, as a result of improved elasticity, compared to BSA-stabilized emulsion. However, clear adhesion remains observed to BSA/RGD, despite slightly lower elasticity, and even unfunctionalized BLG-stabilized interfaces supported some cell proliferation. This may be associated with matrix deposition and remodeling at these interfaces, as cells are able to deposit significant levels of fibronectin fibers at corresponding interfaces. Overall, this study indicates that replacing co-surfactant molecules such as PFBC and achieving bioactivity with readily available scaffold proteins typically encountered in food processing and stem cell technologies is possible.

## 4. Materials and Methods

### Materials and Chemicals

β-lactoglobulin (>90%), BSA (>98%), 1H,1H,2H,2H-Perfluorodecanethiol (97%) and α,α,α-trifluorotoluene (99%) were obtained from Sigma Aldrich Co. The reagent sulfo-succinimidyl 4-(N-maleimidomethyl) cyclohexane-l-carboxylate (sulfo-SMCC 22322) and the SAMSA Fluorescein, 5-((2-(and-3)-S-(acetylmercapto) succinoyl) amino) Fluorescein, mixed isomers (A685) are purchased from Thermofischer Scientific. The fluorinated oil (Novec 7500; dodecafluoro-2-(trifluoromethyl)hexan-3-yl ethyl ether) is from ACOTA. SPR gold coated chips were obtained from Ssens.

### Preparation of emulsions

For the emulsion generation 1mL of fluorinated oil (Novec 7500, ACOTA) containing or not the fluorinated surfactant 2,3,4,5,6-Pentafluorobenzoyl chloride at final concentration of 10 μg/ml and 2mL protein solution (1mg/ml in PBS) were added into a glass vial. The vial was shaken until the emulsion was created and further incubated for 1h at room temperature. The upper liquid phase was aspirated and replaced with PBS 6 times.

### Interfacial shear rheological measurements

For the rheological measurements, a hybrid rheometer (DHR-3) from TA instruments was used with a double wall ring (DWR) geometry and a Delrin trough with a circular channel. The DWR ring has a diamond-shaped cross section that enables the contact with the interface between two liquids to measure the properties. The ring has a radius of 34.5 mm with Platinum-Iridium wires of 1 mm thickness. The Derlin trough was filled with 19 mL of fluorinated oil (with or without surfactant) and, using an axial force procedure. The ring was positioned at the interface by ensuring first contact, followed by lowering by 500 μm from this first contact point, to secure the correct position. After that, 15mL of the PBS solution is placed on the top of the oil phase. Time sweeps were performed at a constant frequency of 0.1 Hz and temperature of 25°C, with a displacement of 1.0 10-3 rad, to follow the protein adsorption at the interface. The protein solution (1 mg/mL) was added after 15 minutes in all cases. Before and after each time sweep, frequency sweeps (with constant displacement of 1.0 10^−3^ rad) were carried out to examine the frequency-dependent behavior of the interface and amplitude sweeps (with constant frequency of 0.1 Hz) to ensure that the chosen displacement was within the linear viscoelastic region ^75^.

### Surface Plasmon Resonance (SPR)

SPR measurements were carried out on a BIACORE X from Biacore AB. SPR chips (SPR-Au 10 × 12 mm, Ssens) were plasma oxidized for five minutes and then incubated in a 5 mM ethanolic solution of 1H,1H,2H,2H-perfluorodecanethiol, overnight at room temperature. This created a model fluorinated monolayer mimicking the fluorophilic properties of Novec 7500. The chips were washed once with water, dried in an air stream and kept dry at room temperature prior to mounting (within a few minutes). Thereafter, the sensor chip was mounted on a plastic support frame and placed in a Biacore protective cassette. The maintenance sensor chip cassette was first placed into the sensor chip port and docked onto the Integrated μ-Fluidic Cartridge (IFC) flow block, prior to priming the system with ethanol. The sample sensor chip cassette was then docked and primed once with PBS. Once the sensor chip had been primed, the signal was allowed to stabilize to a stable baseline, and the protein solution (1 mg/ mL in PBS) was loaded into the IFC sample loop with a micropipette (volume of 50 μL). The sample and buffer flow rates were kept at 10 μL/min throughout. After the injection finished, washing of the surface was carried out in running buffer (PBS) for 10 min. Washing of the surface was allowed to continue for 10 min prior to injection of sulfo-SMCC (at 2.0 mg/mL and volume of 50 μL), at a flow rate of 10 μL/ min. Buffer (PBS) was flown on the sensor chip for 10 min to wash off excess sulfo-SMCC solution and data recording was allowed to continue for a further 10 min. Lastly, RGD solutions (at 1.6 mg/ mL, and volume of 50 μL), at a flow rate of 10 μL/ min were injected and allowed to adsorb for 10 min prior the final washing.

### Circular dichroism

For the circular dichroism measurements transparent emulsions were prepared. For matched refractive index emulsion oil mixture of α,α,α-trifluorotoluene and fluorinated oil were used in order to match the refractive index of water. A 1200 μL volume of fluorinated oil was mixed with 800 μL of α,α,α-trifluorotoluene. The oil mixture was mixed with 2 mL of protein solution at a concentration of 1 mg/ mL. The emulsions were left at room temperature for 1 h, then washed 6 times with PB to remove excess free proteins and used fresh for measurements within 2 h. A 350 μL amount of emulsion was transferred to a sample cuvette for measurement in a Chirascan V100 CD spectrometer. Measurements were carried out at 25°C. α,α,α-trifluorotoluene (99% -547948) was purchased from Sigma Aldrich Co. Measurements were smoothed using the Savitzky-Golay smooth filter with a script written in MATLAB and the secondary composition was estimated using the SELCON algorithm in Dichroweb.

### Bio-functionalization of protein nanosheets

Proteins (BSA, β-lactoglobulin) were dissolved (at a concentration of 1mg/ mL) in PBS (pH 7.4. Emulsions were formed as stated above. The sulfo-SMCC was dissolved in (2 mg/ mL) into 0.5 mL of distilled water under sonication for two minutes and then added to the emulsion for 1 h at room temperature. The upper liquid phase was aspirated and replaced with PBS six times to remove the excess of the sulfo-SMCC. For the SAMSA fluorescein activation, SAMSA-fluorescein (10 mg/mL) was dissolved into 0.1 M NaOH and incubated at room temperature for 15 minutes to remove acetyl protecting groups. 14 μL of 6M HCl were added to 0.2 mL of 0.5 M sodium phosphate at pH 7. To each emulsion, 40 μL of dye solution were added and the upper liquid was replaced again with PBS six times to remove the excess of the dye. After the dye reaction, the emulsion was washed with PBS six times. Samples of each emulsion were transferred to a microwell plate for imaging.

### Mesenchymal stem cell culture and seeding

Mesenchymal stem cells (P3-6, Promocell) were cultured in mesenchymal stem cell growth medium 2 (PromoCell). For proliferation assays, MSCs were harvested with accutase solution containing 0.5 mM EDTA (PromoCell) and incubated at 37°C for 5 min. Cells were resuspended in medium at a ratio 1:1, centrifuged for 5 min at 1200 rpm, counted and resuspended in MSC medium at a desired density. Cells were allowed to adhere and proliferate on these substrates/emulsions in an incubator (37 C and 5 % CO2) for different times points (day three, five and seven of culture), prior the staining and imaging. For the cell spreading assay and matrix deposition assay, cells were seeded at concentration of 10,000 cells per well. For passaging, cells were reseeded at a density of 300,000 cells per T75 flask.

### Human embryonic kidney cells culture and seeding

Human embryonic kidney (HEK293) cells were cultured in DMEM (Thermofisher Scientific) containing 10 % Fetal Bovine Serum (FBS, Labtech) and 1 % Penicillin-Streptavidin (5,000 U/mL). Cells were harvested with trypsin (0.25 %) and versene solutions (Thermo Fisher Scientific, 0.2 g/L EDTA Na_4_ in Phosphate Buffered Saline) at a ratio of 1/9. Cells were resuspended in DMEM at a ratio 1:1 and centrifuged for 5 min at 1200 rpm. HEK293T cells were counted and resuspended in DMEM and seeded onto the substrates at a density of 5,000 cell per well. For passaging, cells were reseeded at a density of 200,000 cell per T75 flask.

### Metabolic assay – CCK8

Cell Counting Kit 8 (CCK-8) assay (Sigma Aldrich - 96992) was used to assess cell viability according to manufacturer’s instructions. Briefly, the CCK-8 reagent was added at each sample and at the standard curve samples (for MSCs – 10k, 25k, 50k, 75k, 100k, 200k, and for HEK293T – 5k, 20k, 50k, 100k, 200k, 300k) and incubated for three hours in the incubator at 37 C. A volume of 100 μL (all samples were triplicates) was taken from each sample and the absorbance was measured at 450nm at day one, three, five and seven for HEK293T and at day three, five and seven for MSCs.

### Immuno-fluorescence staining and antibodies

Samples (emulsions) were washed (dilution and aspiration, followed by addition of solutions) once with PBS and fixed with 4 % paraformaldehyde (Sigma-Aldrich; 8 % for samples in Ibidi well plates) for 10 min at room temperature. Thereafter, samples were washed three times with PBS and permeabilized with 0.2 % Triton X-100 (Sigma-Aldrich; 0.4 % for samples in Ibidi well plates) for 5 min at room temperature. After washing with PBS (three times), samples were blocked for 1 h in 3 % BSA. The blocking buffer was partly removed from the samples, not allowing them to be exposed to air, and the samples were incubated with primary antibodies at 4 °C overnight. Samples were washed six times with PBS and incubated for 1 h with the secondary antibodies (phalloidin (Sigma - P1951), 1:500; DAPI, 1:1000; vinculin (Sigma - V9264), 1:1000; fibronectin (Sigma - F3648), 1:500; Ki67 (Abcam - ab15580), 1:500) in blocking buffer (3% BSA in PBS). After washing with PBS (six times), samples were transferred to Ibidi wells for imaging.

### Immuno-fluorescence microscopy and data analysis

Fluorescence microscopy images were acquired with a Zeiss 710 Confocal Microscope using a 63x and 20x objective to image the MSCs and HEK293T.

### Statistical analysis

Statistical analysis was carried out using OriginPro 9 through one-way ANOVA with Tukey test for posthoc analysis. Significance was determined by * P < 0.05, ** P < 0.01, *** P < 0.001 and n.s., non-significant. A full summary of statistical analysis is provided in the supplementary information.

## Supporting Information

Supporting Information is available from.

## Acknowledgements

The authors are grateful for experimental support and training from Dr Dexu Kong. Funding for this work from the European Research Council (ProLiCell, 772462; ProBioFac, 966740) and a Cyprus Scholarship (IKYK, 841C18) is gratefully acknowledged.

