## Supplementary information for "Engineering of Co-Surfactant-Free Bioactive Protein Nanosheets for the Stabilisation of Bioemulsions Enabling Adherent Cell Expansion"

Alexandra Chrysanthou[1,2], Minerva Bosch-Fortea[1,2] and Julien E. Gautrot[1,2*]

*[1] Institute of Bioengineering and [2]School of Engineering and Materials Science, Queen Mary, University of London, Mile End Road, London, E1 4NS, UK.*

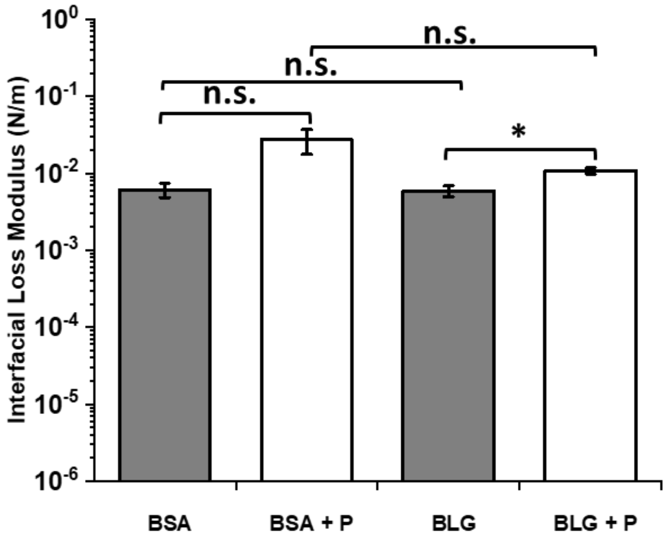

**Supplementary Figure S1.** Interfacial shear loss modulus of BSA, BSA + PFBC, BLG, BLG+PFBC, casein and casein + PFBC at oscillating amplitude $10^{-4}$ rad and frequency of 1 Hz. Error bars are s.e.m; n=3.

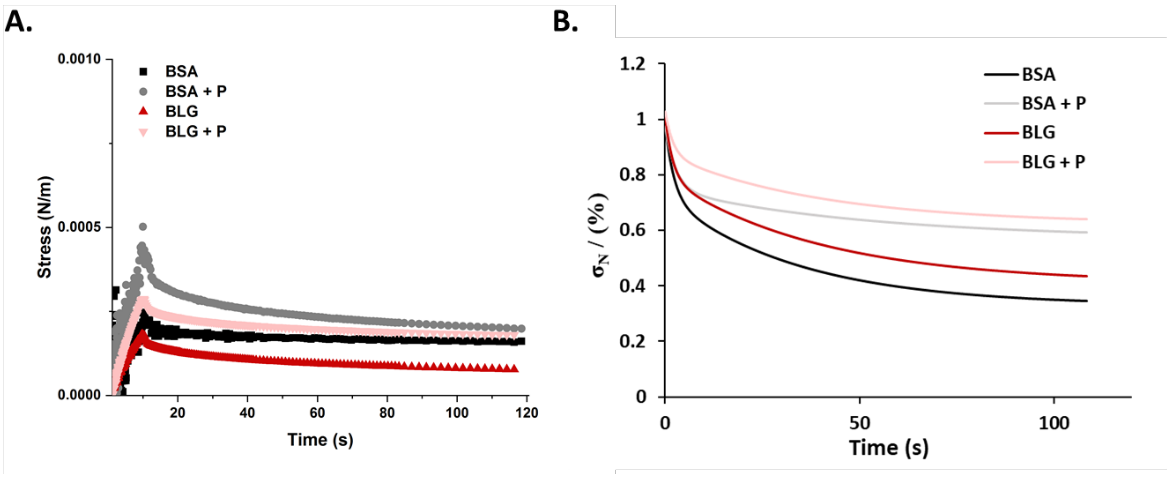

**Supplementary Figure S2.** Stress relaxation experiments carried out on protein nanosheets formed at liquid-liquid interfaces with and without PFBC (10 μg/mL). Data is shown as normalised stress ($\sigma_N$) extracted from stress relaxation experiments at a strain 0.5%.

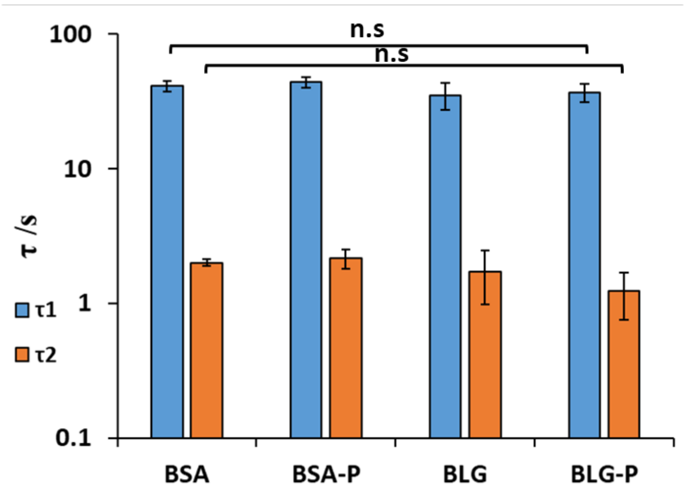

**Supplementary Figure S3.** Characterisation of the stress relaxation profiles associated with protein nanosheets studied. Data were extracted from stress relaxation at a strain of 0.5%. (A) Comparing BSA, BLG and casein (all at 1 mg/mL) with and without PFBC. Error bars are s.e.m; n=3.

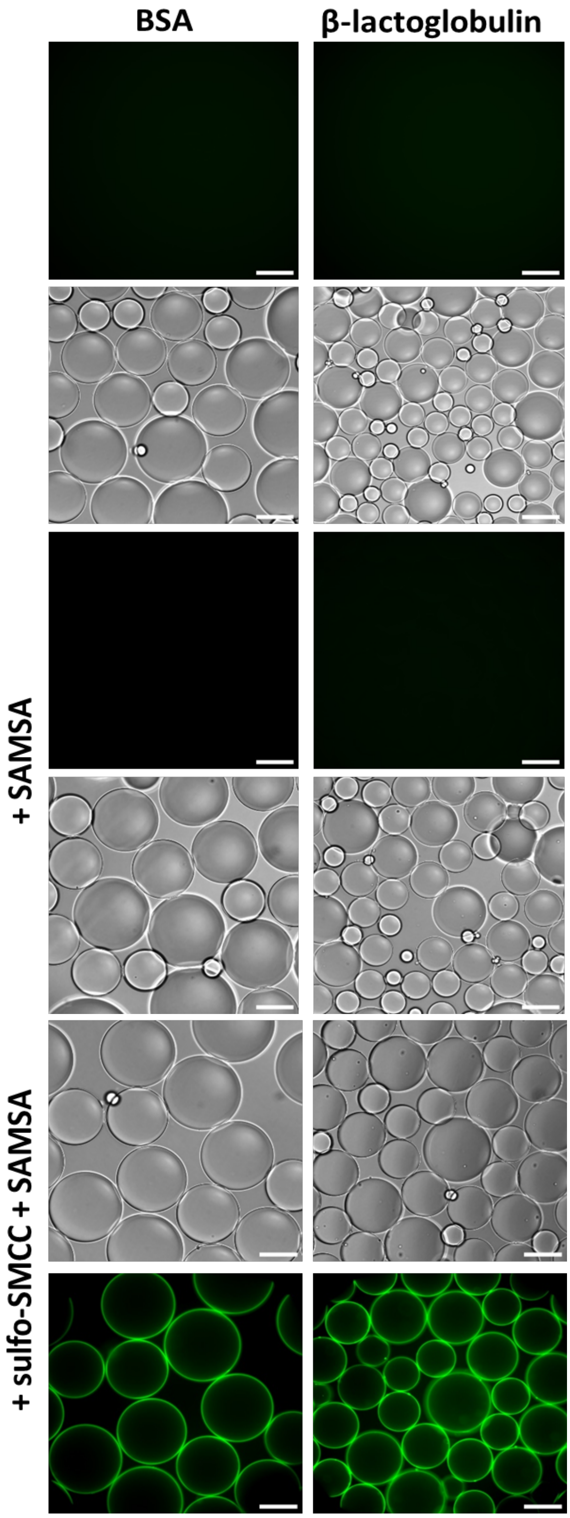

**Supplementary Figure S4.** Epifluorescence microscopy images were SAMSA fluorescein (green, AB F3648) conjugated. Scale bars, 200 μm.

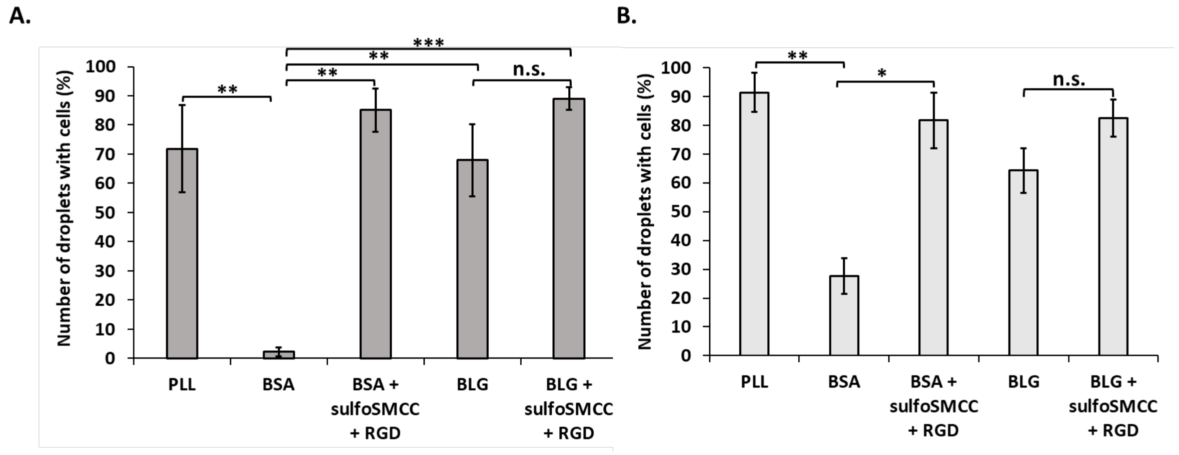

**Supplementary Figure S4.** Percentage number of droplets that were covered with (A) MSCs and (B) HEK293T.

### Supplementary Tables

***Supplementary Table S1*** Summary of statistical analysis of storage modulus data obtained by frequency sweep after the proteins were adsorbed at fluorinated-PBS interfaces with and without co-surfactant (PFBC) at 10 μg/ mL .

| | MeanDiff | Prob | |
|---|---|---|---|
| **BSA + PFBC BSA** | 0.11869 | 0.14564 | **n.s** |
| **BLG + PFBC BLG** | 0.04136 | 0.00209 | ** |
| **Casein + PFBC Casein** | 0.10352 | 0.03876 | * |
| **BLG BSA** | -0.00383 | 0.88433 | **n.s** |
| **Casein BSA** | -0.0356 | 0.0104 | * |
| **Casein BLG** | -0.03177 | 0.01747 | * |
| **BLG + PFBC BSA + PFBC** | -0.08115 | 0.42242 | **n.s** |
| **Casein + PFBC BSA + PFBC** | -0.05077 | 0.69215 | **n.s** |
| **Casein + PFBC BLG + PFBC** | 0.03038 | 0.8718 | **n.s** |

***Supplementary Table S2*** Summary of statistical analysis of storage modulus data obtained by frequency sweep after the proteins were adsorbed at fluorinated-PBS interfaces with and without co-surfactant (PFBC) at 10 μg/ mL.

| | MeanDiff | Prob | |
|---|---|---|---|
| **BSA + PFBC BSA** | 0.0211 | 0.09442 | **n.s** |
| **BLG + PFBC BLG** | 0.00488 | 0.02317 | * |
| **Casein + PFBC Casein** | 0.03089 | 0.01316 | * |
| **BLG BSA** | -2.20E-04 | 0.9853 | **n.s** |
| **Casein BSA** | -0.00564 | 0.01322 | * |
| **Casein BLG** | -0.00542 | 0.01583 | * |
| **BLG + PFBC BSA + PFBC** | -0.01644 | 0.29054 | **n.s** |
| **Casein + PFBC BSA + PFBC** | 0.00415 | 0.90799 | **n.s** |
| **Casein + PFBC BLG + PFBC** | 0.02059 | 0.1718 | **n.s** |

***Supplementary Table S3*** Summary of statistical analysis of data obtained from stress relaxation experiments at a strain of 0.5 % .

| | MeanDiff | Prob | |
|---|---|---|---|
| **BSA-P BSA** | 22.55267 | 0.09127 | **n.s** |
| **BLG-P BLG** | 11.965 | 0.06436 | **n.s** |
| **BLG-P BSA-P** | -2.98533 | 0.9436 | **n.s** |
| **Casein-P BSA-P** | -26.076 | 0.06569 | **n.s** |

| | | | |
|---|---|---|---|
| **Casein-P BLG-P** | -23.0907 | 0.00896 | ** |
| **BLG BSA** | 7.60233 | 0.01709 | * |
| **BLG BSA-P** | -14.9503 | 0.21667 | **n.s** |

***Supplementary Table S4*** Summary of statistical analysis of data of $\tau_1$ values obtained from stress relaxation experiments at a strain of 0.5 %.

| | **MeanDiff** | **Prob** | |
|---|---|---|---|
| **t1 BLG t1 BSA** | 1.48429 | 0.99375 | n.s |
| **t1 BSA-P t1 BSA** | 5.88717 | 0.74876 | n.s |
| **t1 BSA-P t1 BLG** | 4.40288 | 0.87215 | n.s |
| **t1 BLG-P t1 BSA** | 8.50963 | 0.50131 | n.s |
| **t1 BLG-P t1 BLG** | 7.02534 | 0.64101 | n.s |
| **t1 BLG-P t1 BSA-P** | 2.62246 | 0.96783 | n.s |

***Supplementary Table S5*** Summary of statistical analysis of data of $\tau_2$ values obtained from stress relaxation experiments at a strain of 0.5 %.

| | **MeanDiff** | **Prob** | |
|---|---|---|---|
| **t2 BLG t2 BSA** | -0.49612 | 0.84427 | n.s |
| **t2 BSA-P t2 BSA** | 0.76719 | 0.60681 | n.s |
| **t2 BSA-P t2 BLG** | 1.26331 | 0.23655 | n.s |
| **t2 BLG-P t2 BSA** | 0.92303 | 0.46832 | n.s |
| **t2 BLG-P t2 BLG** | 1.41916 | 0.16735 | n.s |
| **t2 BLG-P t2 BSA-P** | 0.15585 | 0.99356 | n.s |

***Supplementary Table S6*** Summary of statistical analysis of data obtained from the SPR data for the protein binding (1 mg/mL) at the perfluorodecanethiol pre-treated chips.

| | **MeanDiff** | **Prob** | |
|---|---|---|---|
| **β-lactoglobulin BSA** | 461.2667 | 0.45494 | **n.s** |
| **β-casein BSA** | 1184.367 | 0.03828 | * |
| **β-casein β-lactoglobulin** | 723.1 | 0.19123 | **n.s** |

***Supplementary Table S7*** Summary of statistical analysis of data obtained from the SPR data for the sulfo-SMCC (2 mg/mL) at the surface of protein layers (BSA, BLG and casein).

| | **MeanDiff** | **Prob** | |
|---|---|---|---|
| **β-lactoglobulin BSA** | 105.0667 | 0.93711 | **n.s** |
| **β-casein BSA** | -80.7333 | 0.9622 | **n.s** |
| **β-casein β-lactoglobulin** | -185.8 | 0.81995 | **n.s** |

***Supplementary Table S8*** Summary of statistical analysis of data obtained from the SPR data for the RGD (1.6 mg/mL) at the surface of the sulfo-SMCC layer.

| | MeanDiff | Prob | |
|---|---|---|---|
| **β-lactoglobulin BSA** | 26.06667 | 0.88869 | **n.s** |
| **β-casein BSA** | 47.4 | 0.68847 | **n.s** |
| **β-casein β-lactoglobulin** | 21.33333 | 0.92352 | **n.s** |

***Supplementary Table S9*** Summary of statistical analysis of data obtained from the epifluorescence images for the SAMSA-flurorescein binding on BSA, BLG and casein emulsion droplets in the presence and absence of sulfo-SMCC.

| | MeanDiff | Prob | |
|---|---|---|---|
| **BLG BSA** | -23.5313 | 0.93412 | **n.s** |
| **Casein BSA** | -37.1652 | 0.8461 | **n.s** |
| **Casein BLG** | -13.6339 | 0.97721 | **n.s** |
| **BLG BSA** | -23.5313 | 0.99966 | **n.s** |
| **Casein BSA** | -37.1652 | 0.997 | **n.s** |
| **Casein BLG** | -13.6339 | 0.99998 | **n.s** |
| **BSA + SAMSA BSA** | 127.9757 | 0.63783 | **n.s** |
| **BSA + SAMSA BLG** | 151.507 | 0.47613 | **n.s** |
| **BSA + SAMSA Casein** | 165.1409 | 0.39065 | **n.s** |
| **BLG + SAMSA BSA** | 6.54822 | 1 | **n.s** |
| **BLG + SAMSA BLG** | 30.07956 | 0.9989 | **n.s** |
| **BLG + SAMSA Casein** | 43.71344 | 0.99363 | **n.s** |
| **BLG + SAMSA BSA + SAMSA** | -121.427 | 0.68331 | **n.s** |
| **Casein+ SAMSA BSA** | 40.45822 | 0.99554 | **n.s** |
| **Casein+ SAMSA BLG** | 63.98956 | 0.96608 | **n.s** |
| **Casein+ SAMSA Casein** | 77.62344 | 0.9267 | **n.s** |
| **Casein+ SAMSA BSA + SAMSA** | -87.5174 | 0.88595 | **n.s** |
| **Casein+ SAMSA BLG + SAMSA** | 33.91 | 0.99805 | **n.s** |

| | | | |
|---|---|---|---|
| **BLG + sulfo-SMCC + SAMSA BSA + sulfo-SMCC + SAMSA** | -649.481 | 0.53284 | **n.s** |
| **Casein+ sulfo-SMCC + SAMSA BSA + sulfo-SMCC + SAMSA** | 96.22956 | 0.98474 | **n.s** |
| **Casein+ sulfo-SMCC + SAMSA BLG + sulfo-SMCC + SAMSA** | 745.7107 | 0.44729 | **n.s** |
| **BSA + sulfo-SMCC + SAMSA BSA** | 4146.55 | 8.79E-04 | *** |
| **BLG + sulfo-SMCC + SAMSA BLG** | 3520.6 | 2.10E-04 | *** |
| **Casein+ sulfo-SMCC + SAMSA Casein** | 4279.945 | 7.43E-04 | *** |

***Supplementary Table S10*** Summary of statistical analysis of data obtained from cell (MSCs) on emulsion droplets after seven days in culture.

| | **MeanDiff** | **Prob** | |
|---|---|---|---|
| **PLL TCP** | -17544.4 | 0.36446 | n.s |
| **BSA TCP** | -75266.7 | 1.52E-05 | *** |
| **BSA PLL** | -57722.2 | 2.16E-04 | *** |
| **BLG TCP** | -30788.9 | 0.03204 | *** |
| **BLG PLL** | -13244.4 | 0.63719 | n.s |
| **BLG BSA** | 44477.78 | 0.00222 | ** |
| **BSA +sulfo-SMCC + RGD TCP** | -29829.6 | 0.03876 | * |
| **BSA +sulfo-SMCC + RGD PLL** | -12285.2 | 0.70135 | n.s |
| **BSA +sulfo-SMCC + RGD BSA** | 45437.04 | 0.00186 | ** |
| **BSA +sulfo-SMCC + RGD BLG** | 959.2593 | 1 | n.s |
| **BLG +sulfo-SMCC + RGD TCP** | -15144.4 | 0.50989 | n.s |
| **BLG +sulfo-SMCC + RGD PLL** | 2400 | 0.99969 | n.s |
| **BLG +sulfo-SMCC + RGD BSA** | 60122.22 | 1.46E-04 | *** |
| **BLG +sulfo-SMCC + RGD BLG** | 15644.44 | 0.47765 | n.s |
| **BLG +sulfo-SMCC + RGD BSA +sulfo-SMCC + RGD** | 14685.19 | 0.54015 | n.s |

***Supplementary Table S11*** Summary of statistical analysis of data obtained from cell (HEK293T) on emulsion droplets after seven days in culture.

| | **MeanDiff** | **Prob** | |
|---|---|---|---|
| **BSA +sulfo-SMCC + RGD BSA** | 284000 | 0.0242 | * |
| **BLG +sulfo-SMCC + RGD BLG** | 297944.4 | 0.02759 | * |
| **BSA +sulfo-SMCC + RGD TCP** | -146056 | 0.25486 | **n.s** |
| **BLG +sulfo-SMCC + RGD TCP** | -148444 | 0.24588 | **n.s** |
| **BLG +sulfo-SMCC + RGD BSA +sulfo-SMCC + RGD** | -2388.89 | 0.99953 | **n.s** |

***Supplementary Table S12*** Summary of statistical analysis of data obtained from the number of droplets covered with cells (MSCs) on emulsion droplets after seven days in culture.

| | **MeanDiff** | **Prob** | |
|---|---|---|---|
| **BSA PLL** | -69.5612 | 0.00242 | ** |
| **BSA + sulfoSMCC + RGD PLL** | 6.0635 | 0.98904 | **n.s** |
| **BSA + sulfoSMCC + RGD BSA** | 75.62472 | 0.00129 | ** |
| **BLG PLL** | -3.88011 | 0.99801 | **n.s** |
| **BLG BSA** | 65.68111 | 0.00367 | ** |
| **BLG BSA + sulfoSMCC + RGD** | -9.94361 | 0.93631 | **n.s** |
| **BLG + sulfoSMCC + RGD PLL** | 17.19101 | 0.689 | **n.s** |
| **BLG + sulfoSMCC + RGD BSA** | 86.75223 | 4.32E-04 | *** |
| **BLG + sulfoSMCC + RGD BSA + sulfoSMCC + RGD** | 11.12751 | 0.90814 | **n.s** |
| **BLG + sulfoSMCC + RGD BLG** | 21.07113 | 0.52247 | **n.s** |

***Supplementary Table S13*** Summary of statistical analysis of data obtained from the number of droplets covered with cells (MSCs) on emulsion droplets after seven days in culture.

| | **MeanDiff** | **Prob** | |
|---|---|---|---|
| **BSA PLL** | -63.759 | 0.00328 | ** |
| **BSA + sulfoSMCC + RGD PLL** | -9.69143 | 0.932 | **n.s** |
| **BSA + sulfoSMCC + RGD BSA** | 54.06757 | 0.01017 | * |
| **BLG PLL** | -27.1195 | 0.26481 | **n.s** |
| **BLG BSA** | 36.63949 | 0.08696 | **n.s** |
| **BLG BSA + sulfoSMCC + RGD** | -17.4281 | 0.64397 | **n.s** |
| **BLG + sulfoSMCC + RGD PLL** | -8.80211 | 0.95079 | **n.s** |
| **BLG + sulfoSMCC + RGD BSA** | 54.9569 | 0.00914 | ** |
| **BLG + sulfoSMCC + RGD BSA + sulfoSMCC + RGD** | 0.88933 | 0.99999 | **n.s** |
| **BLG + sulfoSMCC + RGD BLG** | 18.31741 | 0.60368 | **n.s** |
